# SEACOIN 2.0 – an interactive mining and visualization tool for information retrieval, summarization, and knowledge discovery

**DOI:** 10.1101/206193

**Authors:** Karan Uppal, Eva K. Lee

**Affiliations:** Center for Operations Research in Medicine and HealthCare, Georgia Institute of Technology, Atlanta, GA, 30332.; NSF I/UCRC Center for Health Organization Transformation, Georgia Institute of Technology, Atlanta, GA,30332.; School of Industrial and Systems Engineering, Georgia Institute of Technology, Atlanta, GA, 30332.; 4School of Biology, Georgia Institute of Technology, Atlanta, GA, 30332.; School of Computer Science, Georgia Institute of Technology, Atlanta, GA, 30332.

## Abstract

**Motivation:** The rapidly increasing size of biomedical databases such as MEDLINE requires the use of intelligent data mining methods for information extraction and summarization. Existing biomedical text-mining tools have limited capabilities for inferring topological and network relationships between biomedical terms. Very often too much is returned during summarization leading to information overload.

**Results:** We present herein SEACOIN 2.0, an interactive knowledge discovery and hypothesis generation tool for biomedical literature.SEACOIN generates k-ary relational networks of biomedical terms using a novel term ranking scheme to facilitate efficient information retrieval, summarization, and visual data exploration. Summarization is presented via multiple dynamic visualization panels. We evaluate the system performance in information retrieval and features extraction using the BioCreative 2013 Track 3 learning corpus. An average F-measure of 94% was achieved for document retrieval and an average precision of 88% was achieved for identification of top co-occurrence terms. The system allows interactive mining of complex implicit and explicit relationships among biomedical entities (genes, chemicals, diseases/disorders, mutations, etc.) and provides a framework for hypothesis generation. It also improves our understanding of various biological processes and disease mechanisms.

## 1 INTRODUCTION

Complex diseases such as cancer, diabetes, and cardiomyopathy involve multilevel interactions of cellular processes (Dyugu 2014). Knowledge and discovery of the multitude of interactions and relationships between genes, proteins, metabolites, mutations, epigenetic modifications and environmental factors is essential for understanding the pathophysiology of diseases (Aebersold 2008, Dyugu 2014). Rapid developments in biomedical research over the last two decades have enhanced our ability to study these relationships, which has led to an exponential growth in information available in the form of scientific literature, experimental datasets, and publicly available biomedical databases (Shatkay 2005, Faro 2011).

Currently, over 24 million articles are available in PubMed (http://www.ncbi.nlm.nih.gov/pubmed). In addition to scientific literature, structured and unstructured clinical text information is also recorded in electronic health record systems, which are now being used for phenotyping patients for genetic studies and for epidemic detection (Denny 2013). Studies have shown that it could take up to five years to keep up with the articles that are published per day (Scherf 2005). To further scientific advances, intelligent tools and algorithms are required for rapid processing of the vast amount of free text available in numerous heterogeneous sources.

Biomedical text mining is a growing research area that involves information retrieval, information extraction, named entity recognition, knowledge discovery, and summarization from scientific literature (Rebholz-Schuhmann 2012, Cohen 2013). The concept of literature based discovery was established by Swanson based on his findings of the implicit relationship between fish-oil and Raynaud’s disease (Swanson 1986). Various text-mining tools have been developed since then for inferring gene-gene relationships (Liu 2006), gene-drug interactions (Griffith 2013), drug-disease relationships (Qu 2009), gene-chemical-disease relationships (Davis 2009), gene-disease interactions (Kim 2013), protein-protein interactions (Fernandez 2007, Kwon 2014), drug repurposing (Andronis 2010), and automated hypothesis generation (Liekens 2013). However, the ability to summarize the large body of scientific literature and obtain a systems level overview of the interactions without running into the problem of “information overload” still remains a challenge.

In a recent review, Lu et al. surveyed 28 text-mining tools for searching biomedical literature (Lu 2011). The authors categorized the existing systems based on features such as ranking search results, clustering search results into topics, extracting and displaying semantics and relations, and improving search interface and retrieval experience. Only 2 out of the 28 tools reviewed had graph based visualization capabilities. Visual data mining facilitates exploration of large volumes of data by combining data mining methods with information visualization techniques. Previous studies have shown that the process of data exploration can be made more effective by including the human knowledge in the exploration process (Keim 2002).

Our group has previously developed the SEACOIN system (SEACOIN, Lee 2011), **S**earch **E**xplore Analyze **CO**nnect **IN**spire, as a proof-of-principle tool to address the various needs of biomedical researchers related to literature mining. SEACOIN 1.0 includes features such as depth of retrieved information, familiar and simple-to-use interface, information overload, number of returned pages, and design of result page format that is conducive to users’ understanding of results. Its major summarization features include efficient information retrieval via navigational relationship networks and one-page summarization of abstracts related to a query. The system performance was compared with two other systems, Anne-O-Tate (Smalheiser 2008) and Also Try (Lu 2009); and it was shown that SEACOIN provided most gradual and consistent filtering.

In this paper, we introduce SEACOIN 2.0, which is an improved system that includes several new features including hypothesis generation based on both open discovery and closed discovery models (Andronis 2010) and extractive summarization of top ranked abstracts that are associated with the query. In addition, SEACOIN 2.0 also features technical improvements: a) usage of a controlled vocabulary to limit the k-ary trees to genes, proteins, biological processes, chemicals, species, mutations, microRNAs, histone modifications, diseases and disorders; b) incorporation of text preprocessing steps such as stemming and stop words removal; c) use of point-wise mutual information and tf-idf critieria for establishing query-term association; d) a “history” feature to store previously queried computed co-occurrence networks to reduce the computational time; and e) extractive summarization of the top abstracts.

The key advances of the current research over existing text-mining tools include the usage of k-ary tree structures for query-expansion to enhance information retrieval and visualization of multi-level interactions across biomedical entities in an organized and hierarchical manner. The sub-trees in the network allow users to discover and visualize association of the query term(s) with different biomedical entities, biological processes, environmental exposures, and diseases. The software provides an interactive interface to enhance information retrieval and facilitates discovery of implicit and explicit associations between biomedical entities for hypothesis generation. We demonstrate the utility of the system for information extraction and knowledge discovery using an annotated corpus of 1,112 PubMed abstracts that was used in the BioCreative IV Track 3 task (Arighi 2014). The corpus includes annotations for diseases, genes, proteins, chemicals, and action terms associated with each abstract.

## 2 SYSTEM AND METHODS

SEACOIN2.0 is developed using PHP, Adobe FLEX, Java, MALLET (McCallum 2002), Apache Lucene, and MySQL (Figure 1). The description of the methods and implementation of different components of the system is provided below.

**Fig. 1.**
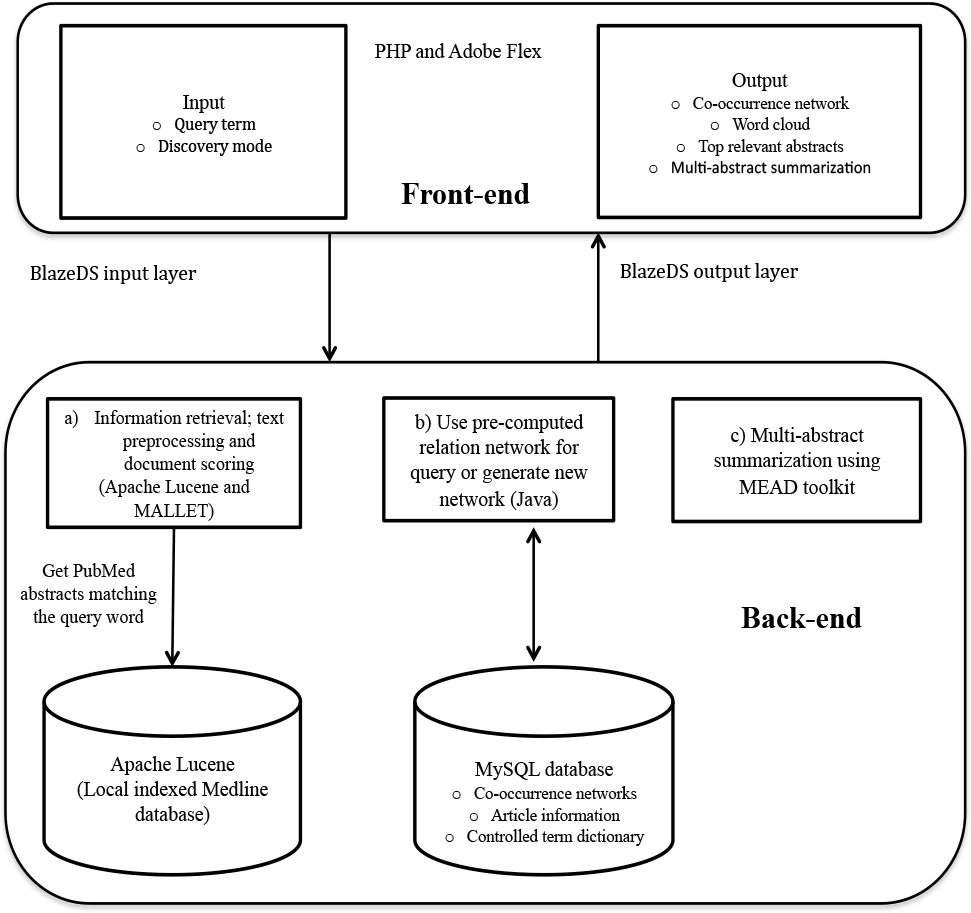
SEACOIN2.0 System Architecture

### 2.1 Medline data

An indexed database of about 14 million abstracts from all articles in the Medline database published between 1975 and 2014 was created using Perl, Entrez e-utilities tools, and Apache Lucene.

### 2.2 Controlled vocabulary dictionary and stop words

First, a term-frequency distribution was generated for ~3.2 million words found in the 14 million abstracts. Each word in this list was then evaluated using databases like MeSH (Coletti 2001), Snomed CT (disorders, findings) and the PubTator database (Wei 2013; a database of includes annotated list of genes, diseases, chemicals, mutations, and species in PubMed), and the following databases in NCBI: Entrez Gene, Protein, PubChem compounds and substances, SNP, Epigenomics, and Taxonomy (NCBI Resource Coordinators 2013).

Additionally, the WordNet database (Miller 1995, Fellbaum 1998) was used to filter words that did not overlap with any terms in PubTator and were not classified under the following categories: noun.state, noun.phenomenon, noun.artifact, noun.process, noun.animal, noun.body, noun.substance, and verb.body.

### 2.3 Text preprocessing

All abstracts were preprocessed in two stages: a) tokenization; b) filtering based on controlled vocabulary and stop words criteria (Figure 2).

#### a) Tokenization

Word tokenization involves segmentation of text into words, punctuations, whitespaces, etc. (Jensen 2006). A dictionary of over 3 million unique words found in the Medline abstracts was generated.

#### b) Filtering based on controlled vocabulary and stop words

A word probability distribution using the entire collection of PubMed abstracts published between 1975 and 2014 was generated. The top most commonly occurring words (p>0.005) along with the stop words used by PubMed (http://www.ncbi.nlm.nih.gov/books/NBK3827/table/pubmedhelp.T43) were removed from the abstracts.

Additionally, words that were not included in the controlled word dictionary (as described in 2.2) were also filtered from the abstracts.

**Fig. 2.**
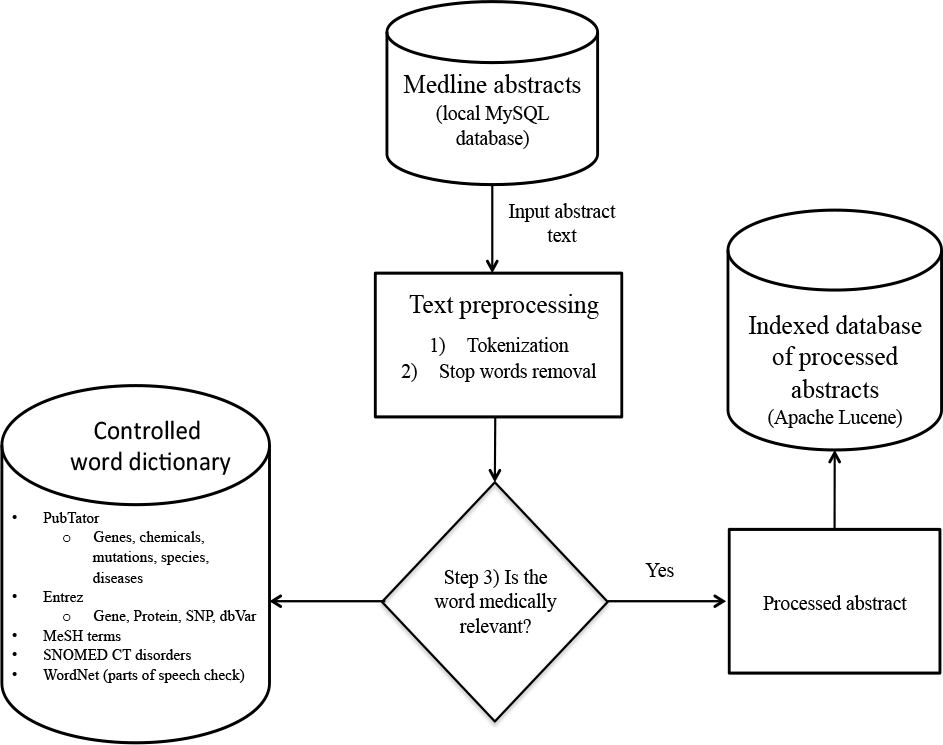
Pre-processing workflow for abstracts

### 2.4 Document indexing, searching, and ranking

In order to facilitate efficient document retrieval during various text-mining stages, the preprocessed PubMed abstracts are stored into an indexed database using the Apache Lucene package (URL: http://lucene.apache.org/). Lucene is an open-source Java based text search-engine library that includes highly efficient search and document scoring algorithms that allow querying based on phrases, wild cards and Boolean operators.

Lucene scores the documents based on the term frequency-inverse document frequency (TF-IDF) scoring scheme as described in the software documentation available at: http://lucene.apache.org/core/3_6_0/api/core/org/apache/lucene/search/Similarity.html. The indexed documents are scored according to their relevance with respect to the query term, t_1_.

### 2.5 Relation discovery and hypothesis generation using k-ary trees

A k-ary tree is a rooted tree where each parent node has up to k child nodes. The maximum number of nodes, N, in a k-ary tree with h levels is given by

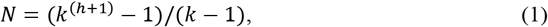

The system generates a 5-ary tree for each query term with h={1,2,3} levels. At each level, the information retrieval process within the system (Section 2.4) is performed in either open or closed discovery mode (Figure 3; Andronis 2010). In the closed discovery mode, abstracts containing all ancestors (e.g.: A and B at level 2 in Figure 3a) in the query term (AB) are used to find the child nodes. In the open discovery mode, the abstracts containing the most immediate predecessor term (e.g.: only term B at level 2 in Figure 3b) in the query are returned. This allows discovery of both implicit and explicit relationships between biomedical entities and generate novel hypothesis as shown in Figure 3 for a 1-ary tree scenario.

**Fig. 3.**
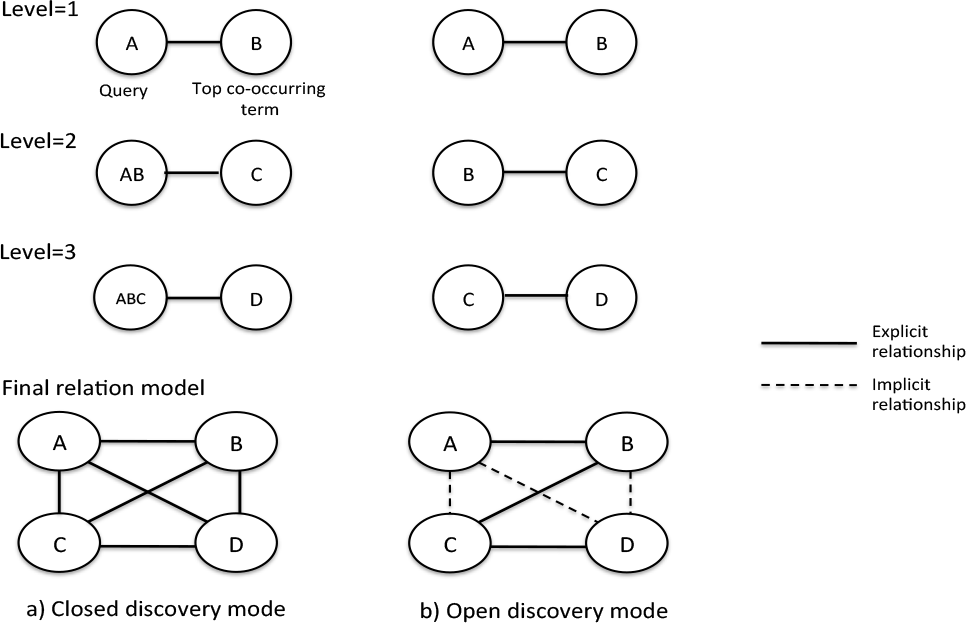
Illustration of the closed (a) and open (b) discovery modes for a 1-ary relation tree with three levels

Users have the option to control the number of levels. A 3-step process is used to generate an ordered tree using the top co-occurring terms at each node (query term) as described below:

Step 1: Term frequency calculation: In this step, a co-occurrence frequency vector for terms co-occurring with the query term, t_1,_ in the indexed abstracts is calculated. The Mallet library (McCallum 2002), a Java-based package for statistical natural language processing, is used to determine the frequency of each co-occurring term, t_2_, in the documents related to the query term, D_t1_. Porter stemmer algorithm (Porter 1980) is used to reduce similar words such as {activated, activates, activating, activation, activators, activator, active, actively} to {activ} in order to determine more accurate term-frequency and co-occurrence estimates. The stemmed words are converted to standard/normal forms once the top co-occurring words have been identified.

Step 2: Next, pointwise mutual information (PMI; Wren 2004, Esteban 2009) between the query term (t_1_) and term t_2_ is computed using the formula in (2). If the probability of association of the two terms is greater than by chance, then a tf-idf criterion is used to assign the association score (3).

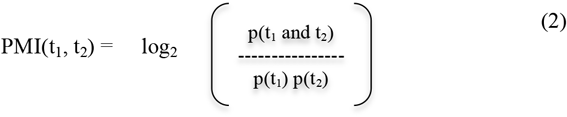

where
p(t_1_) is the probability of term 1 in the corpus, p(t_2_) is the probability of term 2 in the corpus, and p(t_1_ and t_2_) is the probability of co-occurrence of terms 1 and 2 in the corpus.

Step3:

If PMI>1.5, then

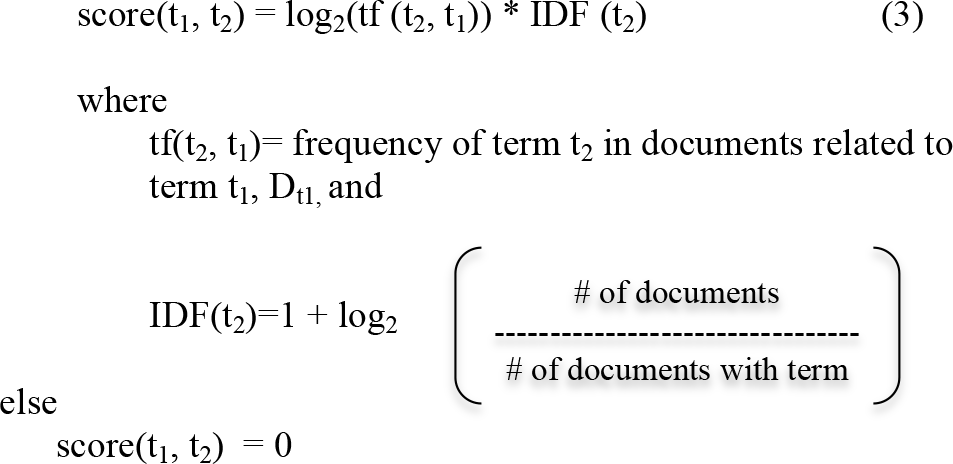

This 3-step approach reduces the risk of incorporating random co-occurrences with commonly used words in the relation networks (Esteban 2009).

Top five co-occurring words are determined for each query based on the PMI criteria and a k-ary co-occurrence network with 3 levels (e.g.: hypertension → angiotensin → pericytes → Nitroarginine) is generated. Interactive word cloud visualization is generated for the top 20 terms to facilitate automated query expansion and document filtering (Lee 2011).

### 2.6 Previously queried computed co-occurrence networks and dynamic database update

SEACOIN 2.0 utilizes MySQL to store previously queried computed networks of MESH terms; it also stores the results for any new queries that are not present in the database. This “memory” feature avoids (repeated) redundant computations and allows the users to retrieve results within 3 seconds for previously submitted queries. For instance, SEACOIN 1.0 took 3 minutes and 30 seconds to compute the word co-occurrence network for “cancer” each time it was submitted as a query. Now the users can retrieve the same network structure within seconds if the query has previously been searched for. SEACOIN 2.0 searches for newly added entries and takes them into account while calculating the word co-occurrence networks at run-time.

### 2.7 Automated multi-abstract extractive summarization

Summarization is the task of representing the information in the original text in fewer words (Cohen 2013). LexRank is a graph-based summarization method that uses cosine similarity and eigenvector centrality to determine the relevance of individual sentences (Erkan 2004). The highest scoring sentence is assigned a score of 1. Our system uses the LexRank algorithm implemented in the PERL MEAD toolkit (http://www.summarization.com/mead/) for identifying most relevant sentences from the top 30 abstracts related to the query. Only the sentences that include at least two words from the controlled vocabulary are included in the summarization process.

### 2.8 MySQL database

The SEACOIN MySQL database includes the PubMed entries (PubMed IDs, Title, Abstracts, Authors, Affiliations, Citation counts in PubMed Central), MeSH entries (MeSH terms and concepts), PubTator terms, stop words, WordNet 2.0 database, and previous queried computed co-occurrence networks (described in Section 2.6).

**Fig. 4.**
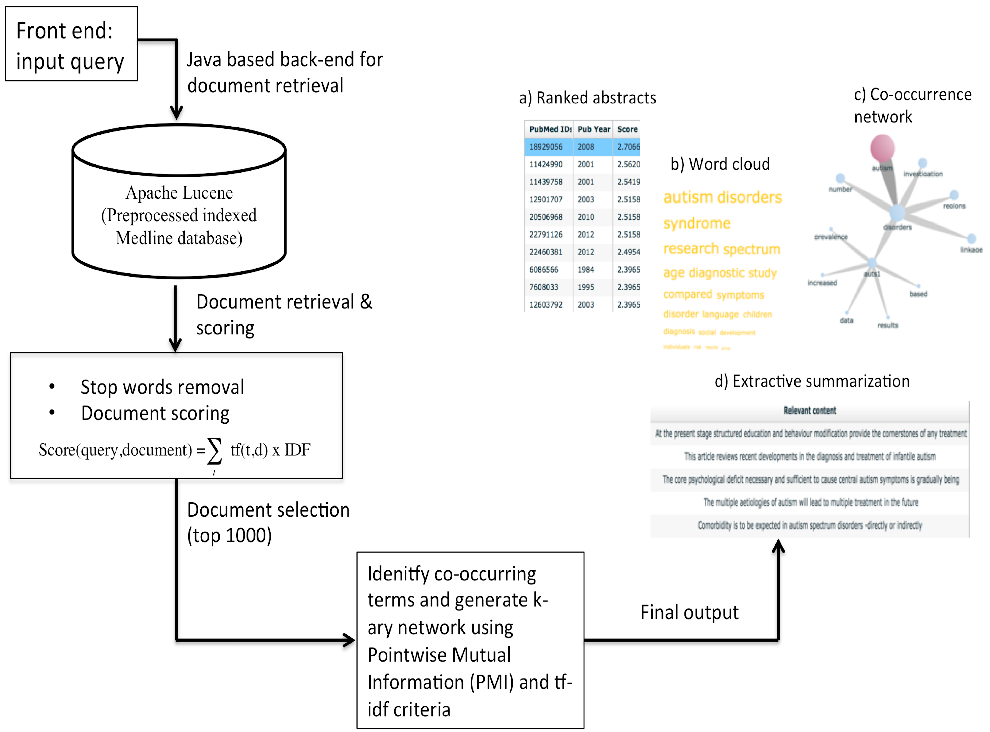
Information flow diagram

These features combined with the features in the original version of SEACOIN can be used to address various scientific queries as illustrated in Section 3.

## 3 EVALUATION AND USAGE

The annotated set of 1,112 abstracts from the BioCreative IV Track 3 CTD learning corpus was used to evaluate the performance of the system using standard text mining metrics such as precision, recall, and balanced F-measure (Cohen 2013). The corpus includes curated annotations for each abstract such as chemical names, gene names, disease names, and action terms (Arighi 2014). Each abstract in the corpus was preprocessed using the workflow outlined in Section 2.2 and indexed using Apache Lucene. Co-occurrence networks were generated as described in Sections 2.3 and 2.4. The following five biomedical terms were used for evaluating the document retrieval and relationship extraction: hypertension, schizophrenia, myocardial infarction, trpv1, and cocaine (Table 1 and 2).

### 3.1 Document retrieval evaluation

Each abstract in the gold-standard corpus was annotated with genes, diseases, and chemicals. On an average, a precision of 89%, a recall of 100%, and a F-measure of 94% is achieved for document retrieval (Table 1).

**Table 1.** SEACOIN 2.0 document retrieval evaluation in terms of precision, recall and F-measure

| \|S\| | Query | Precision | Recall | F-measure |
| --- | --- | --- | --- | --- |
| 1 | Hypertension | 0.897 | 1 | 0.947 |
| 2 | Schizophrenia | 0.727 | 1 | 0.842 |
| 3 | Myocardial Infarction | 0.904 | 1 | 0.95 |
| 4 | trpv1 | 1 | 1 | 1 |
| 5. | Cocaine | 1 | 1 | 1 |
|  | Average | 0.89 | 1 | 0.94 |

### 3.2 Relationship extraction evaluation

An average precision of 88% is achieved for the top 10 co-occurring terms with the query term (Table 2), respectively. The list of terms found by SEACOIN but not present in the curated annotation list include names of enzymes (hce1), physical state (hyperactivity) and anatomical structures (amygdala), demonstrating the ability of the system to detect biologically relevant information in an unsupervised fashion without manual intervention.

**Table 2.**
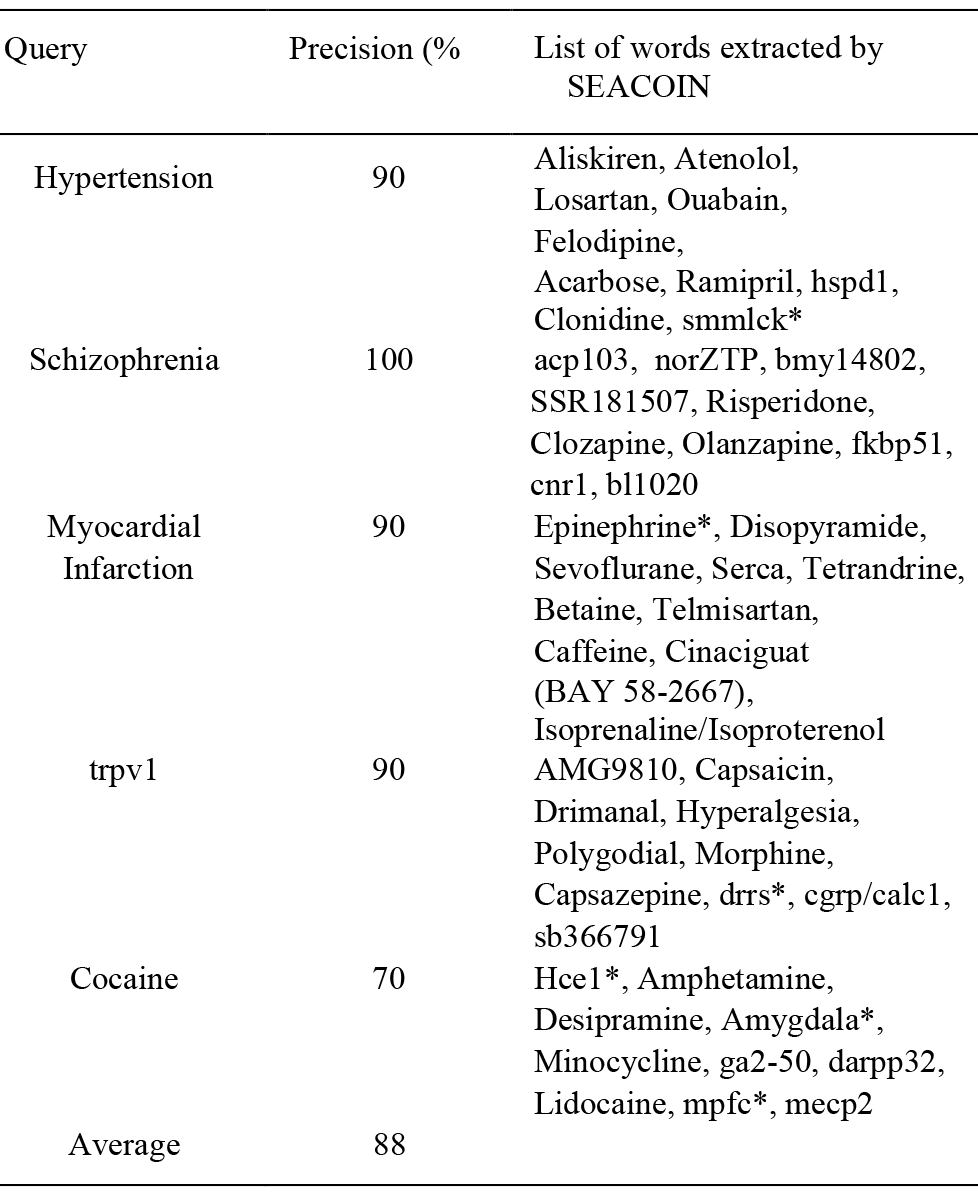
Evaluation of query-term relationship extraction using SEACOIN 2.0 in terms of precision. The [*] symbol represents terms that did not overlap with the curated annotations.

For instance, the search for “schizophrenia” in the evaluation corpus of 1,112 in the open discovery mode identified 11 abstracts related to schizophrenia and one of the subtrees includes schizophrenia (level 0) → bmy14802 (level 1) → dopa (level 2) → fosb (level 3) as shown in Figure 5a. As shown in Figure 5b, “fosb” was not mentioned in any of the abstracts matching “schizophrenia” or “BMY14802”, an anti-psychotic drug. Therefore, it can be hypothesized that the fosb gene and schizophrenia are possibly linked since our knowledgebase is restricted to the abstracts in the evaluation corpus.

An independent PubMed search using “schizophrenia” and “fosb” resulted in 10 hits (at the time of this writing). One of the articles supported the hypothesis that schizophrenia, fosb and BMY14802 are linked. In their recently published work, Dietz et al. showed that the fosb expression levels increased in schizophrenic patients who were prescribed antipsychotic drugs, while no effect in fosb levels was observed in patients who were medication free (Dietz 2014).

**Fig. 5.**
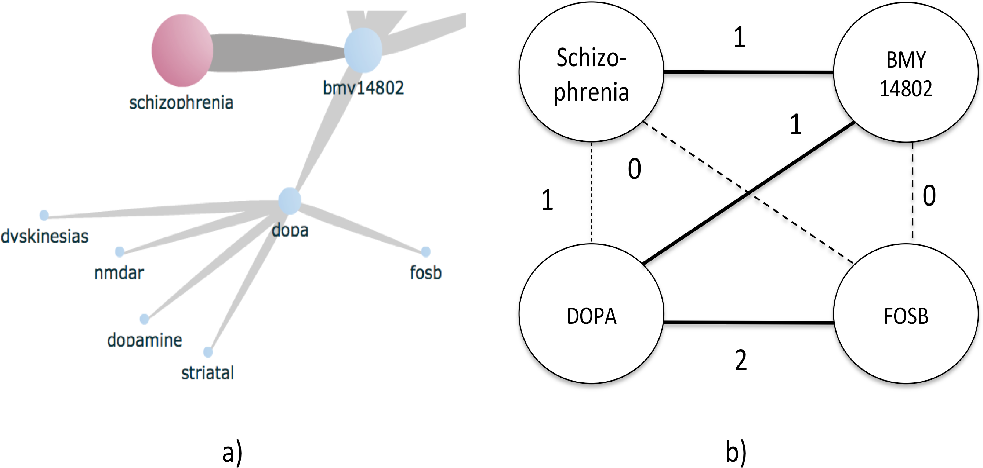
**Fig 5**. Illustration of literature-based discovery based on the evaluation corpus for “schizophrenia” a) A subtree/branch from the relation tree for “schizophrenia” generated by SEACOIN 2.0 shows the relationship between schizophrenia, BMY14802, dopa, and fosb; b) Conceptual representation of the implicit (dotted lines) and explicit (bold lines) relationships in a). The numbers on the edges correspond to the number of abstracts containing the connected terms (nodes).

This example demonstrates the ability of the system to extract implicit relationships from literature to facilitate hypothesis generation and knowledge discovery.

## 4 DISCUSSION

Many tools have been developed that aim to extract important information from biomedical literature. We have previously developed **S**earch **E**xplore **A**nalyze **CO**nnect **IN**spire, SEACOIN, for easy and simple to understand summarization and visualization of medical literature (Lee 2011). SEACOIN is a web-based interactive tool designed using concepts and techniques from text mining, network theory, and visual data mining. It aims to provide better understanding of the biomedical literature by finding associations between biomedical terms and discovering hidden patterns in related/unrelated documents. The early system was designed taking into consideration users’ preferences and concerns related to information overload, simple interface, depth of information control, number of pages returned, and computational time.

SEACOIN 2.0 is an improved version that includes new features including hypothesis generation based on both open discovery and closed discovery models and extractive summarization of top ranked abstracts that are associated with the query. The system also features technical improvements through the use of controlled vocabulary to limit the k-ary trees to biologically relevant terms. It incorporates preprocessing steps including stemming and stop words removal. It also employs point-wise mutual information and tf-idf criteria for defining query-term co-occurrence. To save computational effort, a “history” module is developed to store previously queried computed co-occurrence networks queried by previous users. The new system also includes extractive summarization of the top abstracts.

The system provides a one-page graphical and extractive summarization of the abstracts related to the query. A multi-level interactive topological visualization is used to present the k-ary relation network where each node represents biomedical/clinical terms associated with the query. The open and closed discovery modes for generating the interactive relation networks allow users to discover and explore implicit and explicit relationships between genes, chemicals, proteins, diseases, and mutations as shown in Figure 5. Users can easily traverse the hierarchy of the network by changing the degree of separation. The system allows increase/decrease in the depth of information as users can perform real-time filtering of the network and the corresponding documents by clicking on nodes of interest in the network.

The world cloud is a visual representation of the most frequently co-occurring words with the query term. The terms included in the word cloud are representative of the search term. Users can click on any term in the cloud to update the network and the list of returned documents according to the k-ary string structure. This allows users to dynamically expand their queries and incorporate terms and topics related to the query for enhanced information retrieval. It also provides an alternative and an efficient search strategy to retrieve information as compared to those used by search engines like Google and PubMed.

This flexible structure allows retrieval of large amount of complex information in a simple easy-to understand manner. A table including the list of articles matching the search term is also presented. Users can choose to retrieve more information about each abstract from this table.

The evaluation of document retrieval and co-occurrence network extraction results from SEACOIN2.0 with the BioCreative IV corpus shows that the system can accurately retrieve most relevant documents related to the query without generating too many false positives. It can also discover biologically meaningful query-term associations in both open and closed discovery modes. As demonstrated in Figure 5, SEACOIN 2.0 has the potential to generate novel hypothesis and discover previously unknown connections between diseases, genes, chemicals, etc.

Figure 6 shows the network relationship of glutathione with genes, chemicals, enzymes, exposures, biological processes, and diseases/disorders generated using SEACOIN 2.0 in the closed discovery mode. Glutathione is a tripeptide (cysteine, glutamate, glycine) and is involved in various biological processes such as redox mechanism, immune response, and drug metabolism (Hao 1994, Jones 2002, Chen 2010). It is also known to be associated with various diseases such as liver injury, cancer, cardiovascular disease, age-related macular degeneration, and infectious diseases (Samiec 1998, Jones 2002, Townsend 2003). Additionally, glutathione has been shown to be associated with environmental exposures such as paraquat, smoking, alcohol, arsenic exposure, etc. (Hagen 1986, Yeh 2007, Jones 2011, Hall 2013).

Figure 6a shows the 5-ary relational network for glutathione with 3 levels. The network provides an overview of the associations of glutathione with enzymes (reductase, transferase, peroxidase), diseases (cancer, liver injury, cataract, hypoglycemia, malaria, leishmanisis, hyperhomocysteinemia, ischemia, atherosclerosis, etc.), functions (drug metabolism, involvement in oxidation), organ systems (liver, kidney, etc.), dietary components, drugs (doxorubicin), and environmental exposures (arsenic, alcohol, etc.).

Figure 6b shows level 1 associations of “glutathione AND transferase” that includes kidney, cancer, drug metabolism and anti-cancer drugs like doxorubicin and cisplatin. In the next level, the sub-network for query “glutathione AND transferease AND cancer” is associated with two glutathione transferase genes, gstt1 and gstm1, along with breast, colon, and bladder. Each sub-network represents a different theme providing a systems level coverage of the complex relationships between biomedical entities as compared to commonly used single level search strategies.

**Limitations**: Currently, the system only searches for top five most co-occurring terms with the query and extends the tree up to three levels. Future work will include porting the system to a cloud-computing environment and adopting scalable and distributed computing software such as Apache Hadoop. This will provide a robust computational framework to generate k-ary trees with more nodes and levels. We expect more sophisticated relational theory will also be explored to integrate the large amount of concepts and knowledge into meaningful summary.

## 5 CONCLUSION

SEACOIN 2.0 includes improved methods for document retrieval, information extraction and summarization, and generating co-occurrence networks to discover complex relationships among biomedical terms. We demonstrate herein that SEACOIN 2.0 can be used to retrieve PubMed articles that are most relevant to a query of interest and to generate a collective summary of the previously published studies. This will facilitate discovery of direct and indirect relationships between genes, SNPs, chemicals, and diseases by means of an interactive k-ary word co-occurrence network. Also, the relevance scores assigned to each PubMed article will make the literature review process more efficient as this will limit the focus to a subset of articles of interest rather than thousands of articles, which may or may not be directly relevant to the query term. Hence, the updated SEACOIN system will enhance the ability to extract and summarize knowledge from different articles and help generate hypothesis for future experiments, which is otherwise hard to perform using a simple PubMed search. The ability of the system to extract implicit relationships from literature to facilitate hypothesis generation and knowledge discovery is promising. Further study will be conducted along with biologists in testing some of the potential new hypotheses identified by the system.

**Fig. 6.**
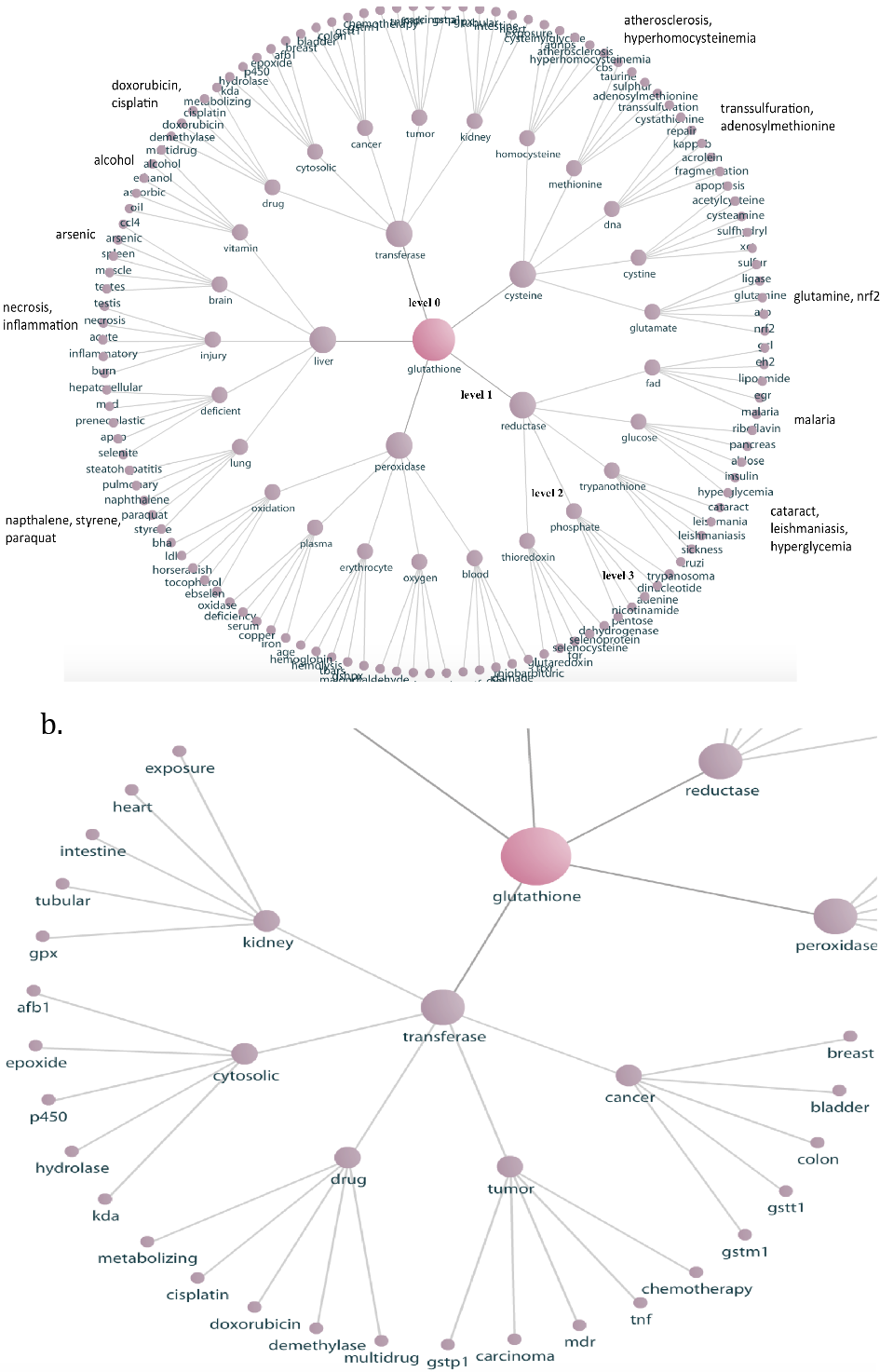
Graph summarization of “glutathione” using SEACOIN2.0. a) The complete three level 5-ary relation network providing an overview of the associations of glutathione with different chemicals, genes, diseases, enzymes, body organs, amino acids, and environmental exposures. b) The 2-level sub-network for query “glutathione AND transferase” that shows its associations with drug metabolism, P450, kidney, and other enzymes; indicates its involvement with multidrug resistance (mdr); and captures the relationship between glutathione transferase, cancer, and genes like gstt1 and gstm1.

## ACKNOWLEDGEMENTS

We would like to thank Dr. Dean P. Jones (Director, Clinical Biomarkers Laboratory, Emory University) for his feedback on the results generated by SEACOIN 2.0.

